# Alpha oscillations reflect suppression of distractors with increased perceptual load

**DOI:** 10.1101/2021.04.13.439637

**Authors:** Tjerk Gutteling, Lonieke Sillekens, Nilli Lavie, Ole Jensen

## Abstract

Attention serves an essential role in cognition and behaviour allowing us to focus on behaviourally-relevant objects while ignoring distraction. Perceptual load theory states that attentional resources are allocated according to the requirements of the task, i.e. its ‘load’. The theory predicts that the resources left to process irrelevant, possibly distracting stimuli, are reduced when the perceptual load is high. However, it remains unclear how this allocation of attentional resources specifically relates to neural excitability and suppression mechanisms. In this magnetoencephalography (MEG) study, we show that brain oscillations in the alpha band (8-13Hz) implement the suppression of distracting objects when the perceptual load is high. In parallel, high load increased neuronal excitability for target objects, as reflected by rapid frequency tagging. We suggest that the allocation of resources in tasks with high perceptual load is implemented by a gain increase for targets, complemented by distractor suppression reflected by alpha band oscillations closing the ‘gate’ for interference.

## Introduction

When people face multiple elements in a scene, selective attention is required to allocate resources in accordance with task demand^1^. *Perceptual load theory* is a highly influential framework arguing that all attentional resources are used to process stimuli within our limited capacity, and therefore whether task-irrevelant distractors are processed depends on the ‘perceptual load’ of task-relevant stimuli^2–4^. Given that visual processing resources are limited, target processing with high load that takes up more resources, will leave fewer resources to process task-irrelevant stimuli. Tasks of low perceptual load, result in a spill over of left over resources to the processing of other – possibly distracting – stimuli. As a consequence, the interference by distracting stimuli will be higher when the load of the target is low, and *vice versa*. In real-life settings, this translates into being unaware of one’s surroundings when engaged in a demanding task. While it is clear that neuronal oscillations are strongly modulated by attention^5^, it remains unclear how these oscillations serve to support the allocation of resources associated with perceptual load theory.

The brain signals most strongly associated with the allocation of attentional resources are neuronal oscillations in the alpha-band (8-13Hz)^6–8^. It remains an open question what drives modulations in the alpha-band. Visual tasks using a simple cue directing attention to one hemifield typically find a hemispheric lateralisation of power in the alpha band over parieto-occipital regions, with decreased power contralateral to the selected object (the target) and a relative increase contralateral to the irrelevant object (the distractor)^6,9–11^. As alpha band oscillations have been shown to reflect functional inhibition^12,13^, the primary role of alpha oscillations is thought to be the suppression of the processing pathway of the irrelevant items as well as engaging the attended pathway by a decrease in alpha power. Perceptual load theory predicts a reduction of resources for distractors with increased target processing load^14^. However, it remains unclear how the theory relates to modulations in the alpha band. In fact, as more salient distractors increase the need for suppression^15,16^, alpha band power increases may also be driven by the nature of the distractor. While some studies have shown support for a modulation of alpha band power contralateral to anticipated distractors^11,17–20^, other studies question the role of alpha oscillations in suppression of distractors during spatial attention^21^. The pattern of alpha band activity tracks the spatial deployment of attention^22^, but this pattern does not seem to be consistently driven by the anticipated location of distractors^23,24^. Furthermore, a body of behavioural studies have concluded that flexible allocation of distractor suppression is minimal or absent, and points to a slower, spatially fixed, mechanism that is learnt with experience^25–27^. This has led to the proposal that suppression of distractors and the facilitation of targets rely on different mechanisms^23^.

It remains unknown how target facilitation and distractor suppression are implemented at the neuronal level. In the past, alpha-band oscillations have been linked to gain control^12,28^, where states of high alpha band power are associated with low cortical excitability^29,30^. However, recent evidence from MEG using rapid frequency tagging suggests that this relation may not hold when considering trial-by-trial fluctuations^31^. Similarly, in a recent EEG study^32^, early visual gain, as indexed by the steady-state visual evoked potential (SSVEP), did not show any dependency on fluctuations in alpha-band power in a selective attention task. These findings were complemented by an EEG study^33^ reporting increased target processing, as indexed by SSVEP, but no reduction in distractor processing, despite increased alpha band power contralateral to a distractor. This questions a direct link between alpha oscillations and stimulus gain; rather alpha oscillations might implement a gating mechanism just downstream to early sensory regions^31^.

In the current MEG study, we investigate target facilitation and distractor suppression in the context of perceptual load and the role of alpha-band oscillations in sensory gain control versus down-stream gating. By using a cued spatial attention task manipulating target load as well as distractor saliency, we aim to uncover how target load and distractor saliency modulate posterior alpha oscillations as well as neuronal excitability. To assess neuronal excitability we use *Rapid Frequency Tagging* (‘RFT’^34^) on both the target and distractor stimuli. The use of MEG allows for localising the neuronal sources of alpha oscillations and the RFT signal.

In line with perceptual load theory, we hypothesised that if increased load of the target drives the allocation of spatial attention, this would be reflected by an increase in alpha band power contralateral to the distractor. Likewise, we expect RFT power, reflecting neuronal excitability, to be reduced contralateral to the distractor in the case of high target load^35–37^. We also hypothesised that, if spatial attention reflects the suppression of anticipated distractors, the salience of the distractor is a driving factor. Increased distractor saliency should result in increased alpha band power and decreased in RFT power contralateral to the distractor. These hypotheses, pre-registered at https://osf.io/ha4vw/, are not mutually exclusive *per se*. As an additional point of exploration not covered in the pre-registration, recent findings have highlighted the possibility that alpha-band oscillations do not implement gain modulation but rather gating in down-stream regions^31^. We therefore also set out to investigate that if alpha band oscillations implement stimulus gain, RFT power should show the inverse pattern of modulations of alpha oscillatory power. If, however, alpha implements gating at a different stage, these modulations may be independent.

## Results

Subjects (N=30) performed a spatial attention task in which a cue directed the attention to a face in either the left or right hemifield (Fig. 1), while recording MEG. The task was to discriminate a subtle change in the attend face (eye-movement direction) after a delay of unpredictable duration. Noise was added to the face stimuli to increase the perceptual load (due to reduced visibility of the stimulus eye movement) of attended target processing. Noise was also added to the distractor stimulus, independently of the target, thus manipulating the distractor conspicuousness of eye movement direction, which we henceforth refer to as salience (Fig. 1C). Reaction times were recorded and sorted according to the target load condition and distractor salience.

**Figure 1.**
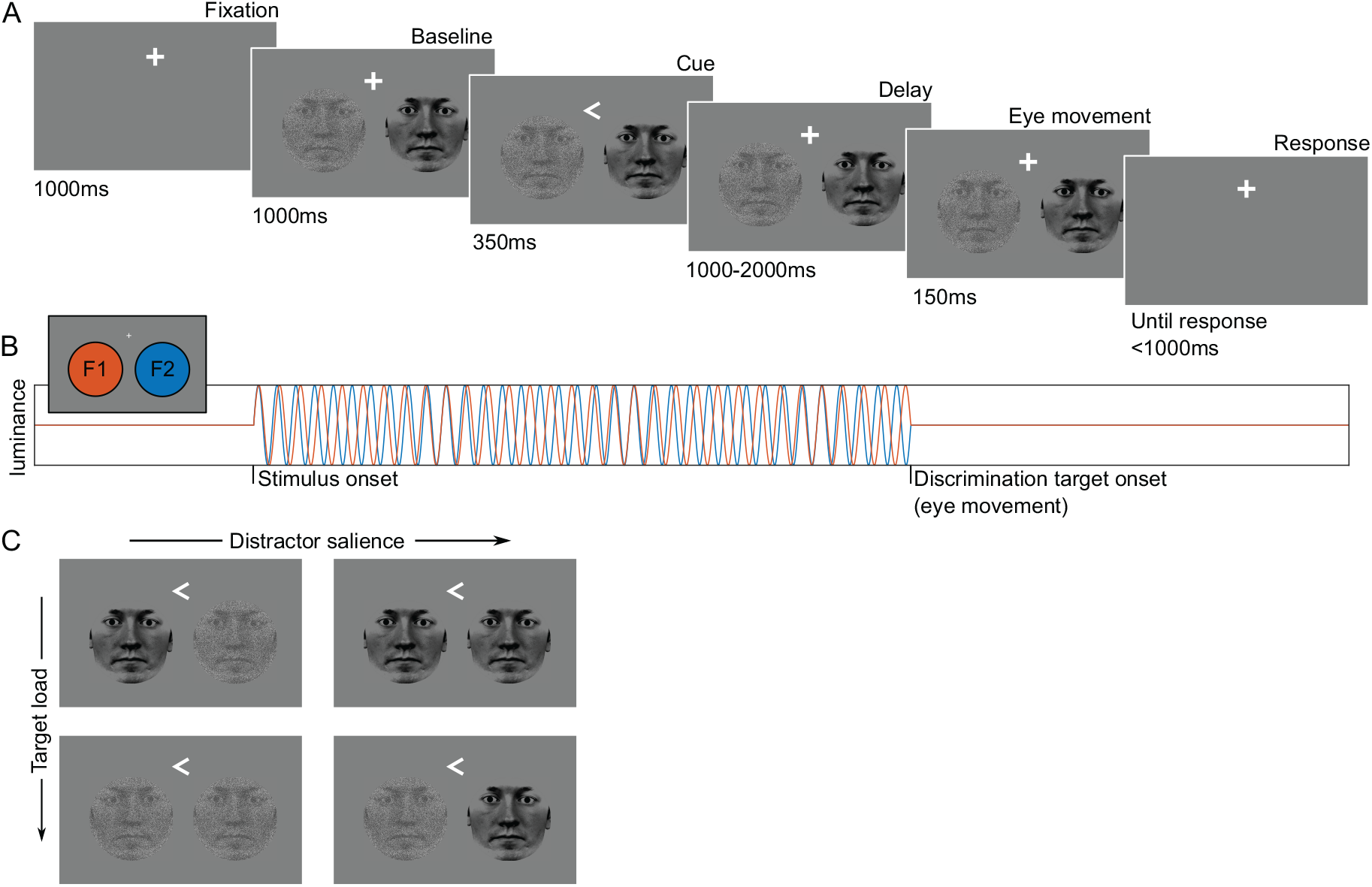
Overview of the cued discrimination task. A) After fixation, subjects were presented with two face stimuli. A directional cue indicated the target stimulus. After a variable delay, a small eye movement occurred in the face stimuli. Subjects indicated the direction of the eye movement by button press. B) The luminance of the white portions of the stimuli flickered at 63 and 70Hz (rapid frequency tagging; RFT), from stimulus onset until the eye movement change of the stimuli. C) The cued target stimuli were masked with noise or not (respectively high versus low perceptual target load). Likewise, the uncued distractors were masked with noise or not (respectively low versus high salient distractors). Noise levels are increased for illustration purposes and stimulus sizes are not to scale.

### Behaviour

The behavioural effect of the perceptual load was reflected in the subjects’ reaction times during the task (Fig. 2A). A repeated measures ANOVA on the reaction times with factors ‘target load’ and ‘distractor salience’ revealed a significant main effect of ‘target load’ (*F_(1,29)_* = 132.1, *p* < .001, *partial η^2^* = .82), demonstrating that the participants responded slower to high compared to low target loads, as would be expected with a higher level of noise on the target stimulus. There was also a main effect of ‘distractor salience’ (*F_(1,29)_*=6.7, *p* = .015, *partial η^2^* = .19) meaning that participants responded slower to the target in the presence of a salient compared to a noisy distractor. This shows that the salient distractors were indeed effective in interfering with performance. Critically, a significant interaction was found between ‘target load’ and ‘distractor salience’ (*F_(1,29)_* = 12.9, *p*=.001, *partial η^2^* = .31). This is explained by slower target-responses in the presence of salient compared to noisy distractors, but only when the target load was low (Fig. 2B). When the target load was high, we observed no effect of distractor salience. Thus, distractor interference was eliminated in conditions of high target load, as predicted by perceptual load theory.

**Figure 2.**
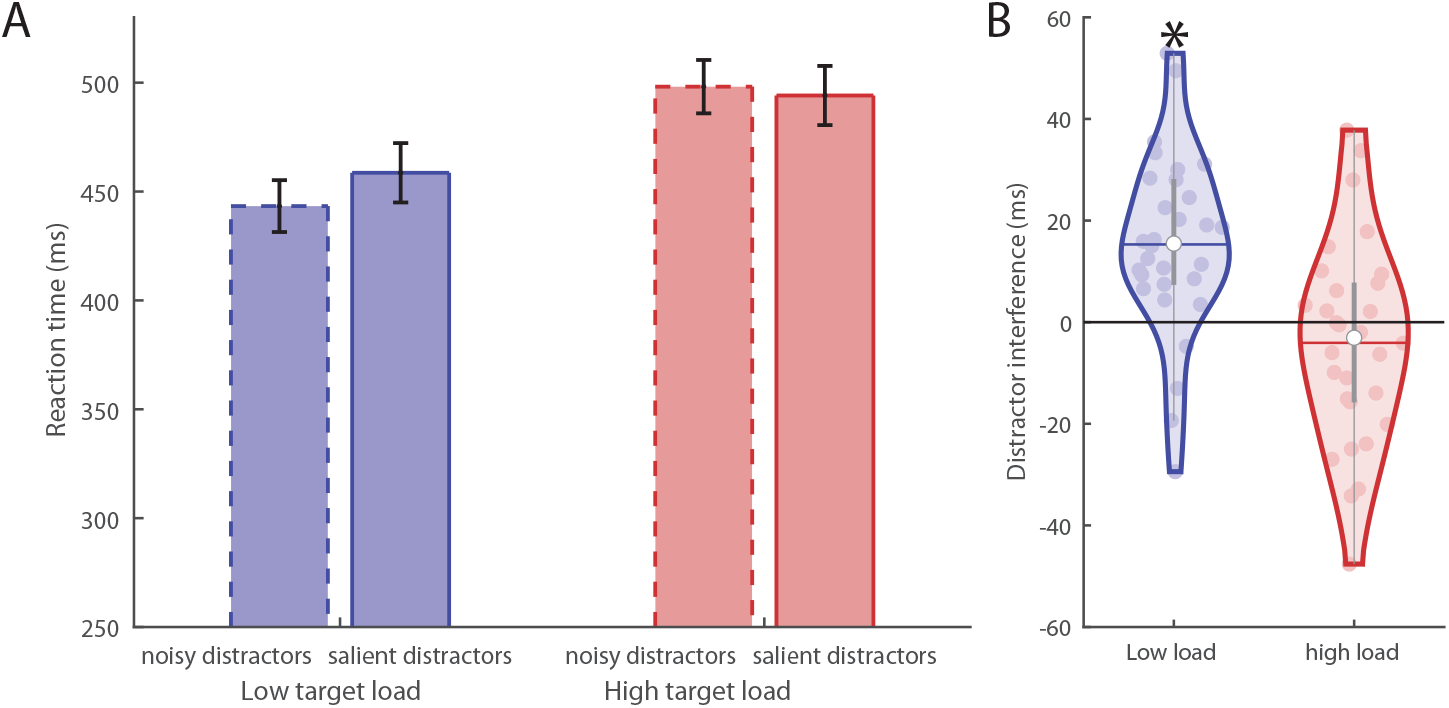
Behavioural results (reaction times to targets) of the discrimination task. (A) When considered per condition, participants responded slower to high (red) compared to lower (blue) target loads. (B) Distractor interference (salient compared to noisy distractors) was significant when the target load was low (M_low_ = 15.3ms, M_high_ = −4.07ms, paired samples t-test t = 3.6, p < .001), but not when the target load was high (p > .25). These findings are consistent with perceptual load theory.

### Effects of spatial cuing

#### Alpha band power

To test the effects of directed attention and select sensors for the subsequent analysis, we compared trials where attention was cued to the left with trials cued to the right hemifield (Fig. 3A). The time-frequency representations of power modulations as well as the topographic representation show a decrease in alpha power contralateral to the target accompanied by an increase contralateral to the distractor. The marked locations in the topographic plot indicate the selected sensors of interest that showed the largest attentional modulation. Figure 3B shows the time-course of the alpha power relative to a pre-cue baseline, locked to the cue (left), and discrimination target onset (right). Importantly, the alpha power increases contralateral to the distractor, but not the target.

**Figure 3.**
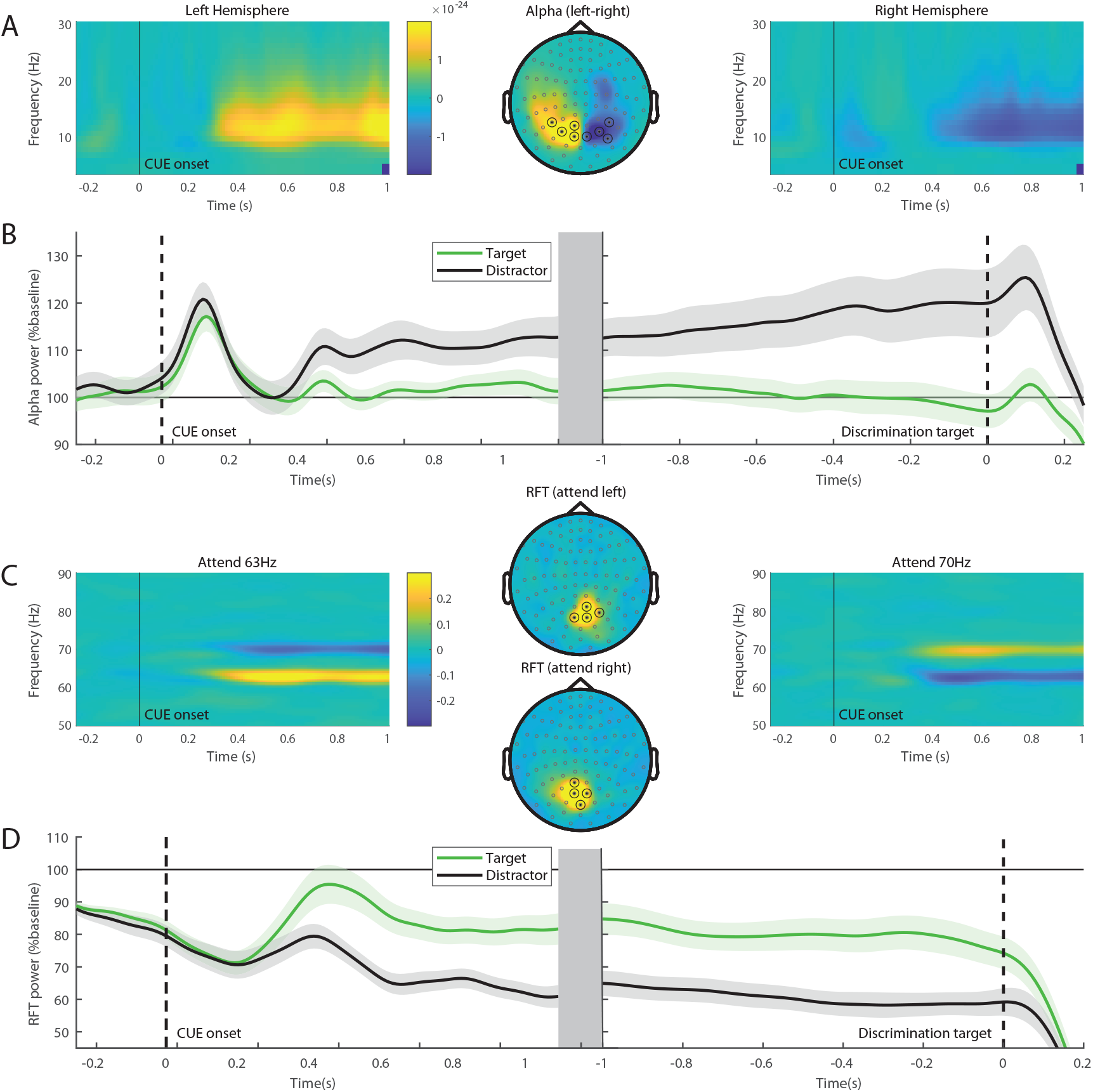
Attentional modulation and sensor selection. A) Alpha band (8-13Hz) modulations were quantified by considering the left versus right directional cues. The modulations in time-frequency representation of power showed a sustained alpha power increase ipsi-laterally to the cued hemifield (left) and a power decrease contralaterally (right). The topographic map demonstrated a power modulation over parieto-occipital areas; the marked location indicates the sensors used in the subsequent analyses. B) Time-courses of the alpha band power relative to pre-cue baseline contralateral to target (green) and distractor (black). We show this with respect to Cue onset (left) and discrimination target onset (right). C) Rapid Frequency Tagging (RFT) power modulations. The time-frequency representations show an increase in power contralateral to the target and a decrease contralateral to the distractor. The target was flickering at 63Hz (left) and 70Hz (right). The topographic plot (centre) shows these modulations to be over central occipital areas. The marked location indicates the sensors used in the subsequent analyses. D) Time-courses of the RFT power relative to pre-cue baseline for target frequencies (green) and distractor frequencies (black) with respect to Cue onset (left) and discrimination target onset (right).

#### Rapid frequency tagging (RFT)

Both target and distractor stimuli flickered at either 63 or 70Hz, i.e. were ‘frequency tagged’ (Fig. 1B). As can be seen in figure 3C-D, RFT power related to the target shows a sustained increase after cue onset relative to distractor power, indicating that RFT power is sensitive to attentional modulations. Unlike alpha band power, attentional selection is reflected in increased RFT power and can thus be considered a proxy for neuronal excitability associated with attentional gain. Note that we found a generic decrease of RFT power over time, regardless of condition or attention direction, which is why the RFT power after cue onset is generally below baseline. Next we examine the effects of perceptual load and distractor salience on attentional modulations in the alpha band and RFT power.

### Effects of perceptual load and distractor salience

As shown, by increasing the load of the target, distractor interference is abolished (see Fig. 2B). Here we relate this to neural suppression (as reflected by alpha amplitude) and excitability (as measured with rapid frequency tagging amplitude). For the alpha band we hypothesise that attentional modulations may be implemented by either a stronger suppression of the distractor with high target load, as reflected by increased alpha band activity contralateral to the *distractor*, or increased resources allocated to the target, as reflected by decreased alpha band power contralateral to the *target*, or both. Rapid frequency tagging (RFT) power, reflecting neuronal excitability or gain, is expected to show the opposite pattern to alpha band power, with target load increases being associated with decreased RFT related to distractor processing, increased RFT related to target processing or both. We also tested the hypothesis that distractor salience drives the distractor suppression effects.

#### Alpha band

##### Alpha band power contralateral to the target

An ANOVA was performed on target-related alpha band power 500-1350ms after cue offset with factors ‘target load’ and ‘distractor salience’. The first 500ms post-cue were discarded to avoid the evoked response of the cue. There were no significant main effects of ‘target load’ or ‘distractor salience’ (Fig. 4A, left) and no interaction (all F_(1,29)_ < 2.44, all p > .12), suggesting neither perceptual load, nor distractor salience affects alpha band power related to the target stimulus.

**Figure 4.**
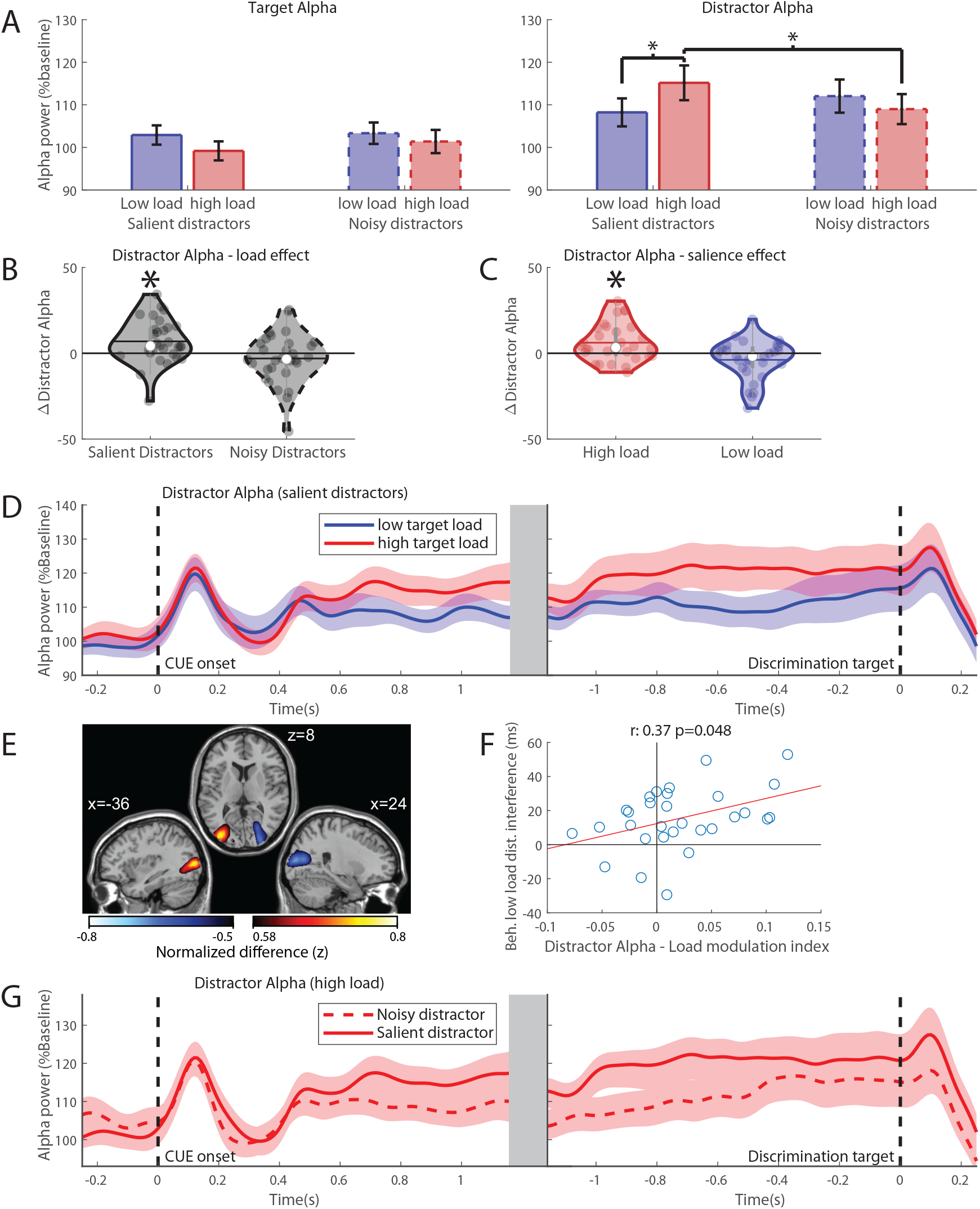
Effects of target load and distractor salience in the alpha band. A) Average alpha band power per condition, contralateral to target (left) and distractor (right) for high (red) and low (blue) target load conditions with noisy (dashed) or salient (solid) distractors 500 – 1350 ms after cue onset. Error bars denote the standard error of the mean (SEM). B) The effect of target load (high-low target load) for salient and noisy distractors. The mean is denoted by the horizontal line, the median is denoted by the white dot and interquartile range by the grey vertical bar. C) The effect of distractor salience (high-low salience) with high and low target load. D) Distractor-alpha power time courses for high (red) and low (blue) target load locked to cue onset (left) and discrimination target onset (right). Importantly the distractor-alpha was larger for high compared to low target-loads, consistent with the hypothesis that alpha power implements the distractor suppression according to perceptual load theory. E) Source modeling of the distractor-alpha modulation of high versus low target loads (cue-locked, 5 – 1.35 s) collapsed over left and right attention displayed as attention to the left (left hemisphere is contralateral to distractor). Plot shows top and bottom 1% values in the grid-points. F) Correlation between average load modulation index (distractor alpha (high load - low load) / high load + low load), cue-locked, 500 – 1350ms after cue onset) of distractor alpha and behavioural distractor interference (RT_noisy_ – RT_salient_) under low target load. This shows that the larger the difference in distractor alpha power (i.e. lower alpha power with low load), the larger the behavioural distractor interference. G) Distractor alpha power time courses for salient and noisy distractors relative to cue onset (left) and discrimination target onset (right) under high target load.

##### Alpha band power contralateral to the distractor

The same 2×2 ANOVA applied to the alpha band power contralateral to the distractor revealed a significant interaction between ‘target load’ and ‘distractor salience’ for alpha band power contralateral to the *distractor* (*F_(1,29)_* = 12.6, *p* = .001, *partial η^2^* = .30) (see Fig. 4A, right). We hypothesized that the effect may be driven by the load of the target. Indeed, post-hoc tests reveal a significant increase in alpha amplitude with increased target load when the distractors were salient (paired samples t-test, *t* = −2.97, *p* = .006, significant with Bonferroni corrected α = .0125) but not when distractors were noisy (paired samples t-test, *t* = 1.14, *p* = .27), see figure 4B and 4D for this effect over time. This shows that perceptual load of the target indeed drives alpha band increases contralateral to the distractor, given that the distractor is sufficiently salient. Source modelling (Fig. 4E) places this load-driven alpha band increase in the extrastriate cortical areas (peak MNI coordinate: x:-36 y:-84 z:8, BA 18/19), extending into fusiform gyrus (BA 37). Here the warm coloured source denotes the increase in alpha power (high versus low targets) contralateral to the distractor. Next, we asked if the target load-driven modulation in distractor-alpha power predicted behaviour. To this end, we consider behavioural interference, i.e. the increase in reaction time caused by an increase in distractor salience relative to the modulation of alpha band power contralateral to the distractor. The modulation of distractor alpha power due to increased target load shows a significant positive correlation (*r*_spearman_ = .37, *p* = .048) with the behavioural interference caused by the distractor (Fig. 4F). This positive correlation demonstrates that subjects with larger distractor interference effects in the low target load conditions had a larger difference in distractor-alpha between high and low load. Thus a greater increase in alpha power contralateral to the distractor with high perceptual load may be required to suppress a more potent distractor that produces larger behavioural interference in conditions of low perceptual load; this is also consistent with the target load effects found on alpha power only for the salient (but not for the noise masked) distractors.

Our results show that attentional modulations are driven by the load of the target. However, as hypothesised and evident from the omnibus ANOVA, the salience of the distractor can drive alpha band modulations. When target load is high, post-hoc paired samples tests show that alpha band power for the distractor is significantly increased when distractors are salient relative to noisy (paired samples t-test, *t* = 3.13, *p* = .004 < α = .0125). This effect, however, does not occur with low target load (paired samples t-test, *t* = −1.81, *p* = .081 > α = .0125), see figure 4C. Figure 4G shows the time-course of alpha band power contralateral to distractors with high target load. A clear increase in alpha band power is observed when distractors are salient, but only when target load is high.

However, these modulations of alpha power do not correlate with the observed behavioural effect (distractor interference, *r*_spearman_ = 0.07, *p* = .72). This suggests that the difference in alpha power between noisy and salient distractors, although significant, is of limited behaviourally relevance in the light of our paradigm. This may be due to the inability of the noisy distractor to cause distractor interference, making the level of associated alpha power, and suppression, less relevant.

In summary, when perceptual load increases, alpha band power contralateral to the distractor increases, reducing behavioral distractor interference seen with low perceptual load. The salience of the distractor is important, as this perceptual load effect only occurs with salient distractors. High salience of the distractor also increases alpha band power associated with the distractor. However, the alpha band power increase due to perceptual load correlates with behavior, whereas the distractor salience does not. This indicates that perceptual load may be the key driving factor in alpha band modulations related to the distractor, whereas sufficient distractor salience is a pre-condition.

#### Rapid frequency tagging

##### RFT target power associated with the target

As for the alpha band, an ANOVA was performed on the rapid frequency tagging (RFT) power averaged in the 500 – 1350ms interval after cue onset, over all conditions. Analysis of RFT-power associated with the *target* (see Fig. 5A), revealed a significant *main effect* of ‘target load’ (*F_(1,29)_* = 27.3, *p* < .001, *partial η^2^* = .49), where overall RFT power is stronger with high (M = 0.87, *SEM* = 0.046) than low target load (M = 0.80, *SEM* = 0.043). Additionally, there was a significant *interaction* between ‘target load’ and ‘distractor salience’ (*F_(1,29)_* = 4.44, *p* = .044, *partial η^2^* = .13) reflecting a stronger effect of target load in the presence of salient distractors (paired samples t-test, *t* = −7.83, *p* < .001 < α = .0125 Bonferroni corrected) compared to noisy distractors (paired samples t-test, *t* = −2.37, *p* < .025 > α = .0125), see figure 5B. Thus, with salient distractors, target RFT power with high load is higher than under low load, as can be seen in figure 5A and B. This shows that neuronal excitability associated with the target is increased with target load. Alternatively, this may point to decreased target processing due to distractor interference when load is low and distractors are salient. The time-course of target RFT for high and low target load, shown in figure 5C, clearly shows a higher RFT power with high target load, which is especially pronounced shortly following cue offset (0.35s). Source modelling reveals this increase to be very focal and situated in primary visual cortex (peak MNI coordinate: x:8, y:-90, z:6, BA 17), extending only slightly beyond (BA18), see figure 5D. Note that, due to the very focal nature of the source, the top 1% values may contain artifactual clusters, such as the frontal cluster. While not significant, there is a trend towards a negative correlation between the target RFT distractor effect (i.e. the difference in target power with salient vs. noisy distractors) and the behavioural distractor interference, with low target load which is clearly absent with high target load (Fig. 5E). This may suggest that the target RFT power is reduced when salient distractors are present when the target load is low, but the distractors have no effect when the target load is high, as predicted by perceptual load theory.

**Figure 5.**
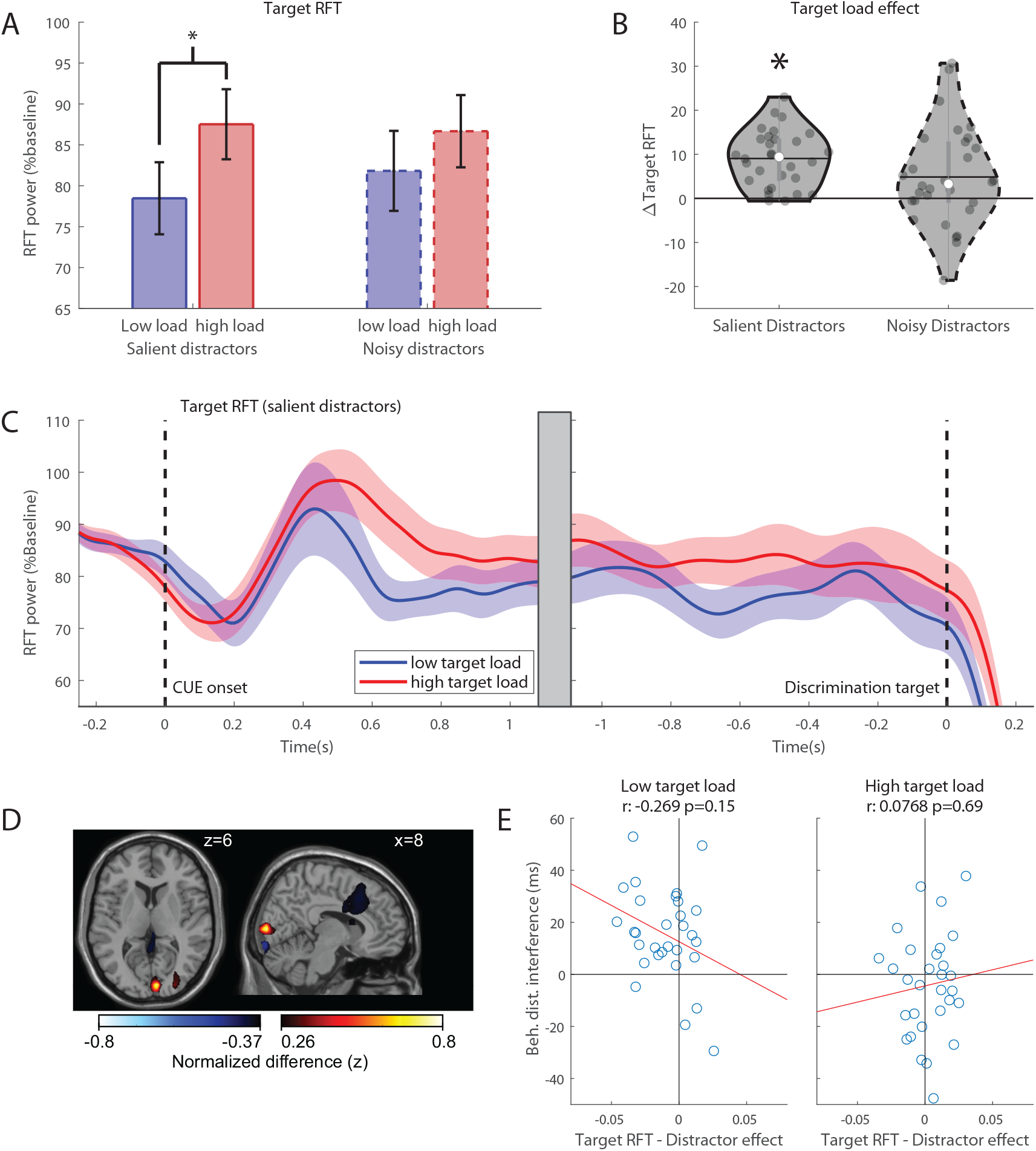
Load effects in RFT power. A) Target RFT power per condition. Average RFT power related to the frequencies of the target for high (red) and low (blue) target load conditions with noisy (dashed) or salient (solid) distractors 500 – 1350ms after cue onset. Error bars denote standard error of the mean (SEM). B) Highlight of the significant effect of target load in target RFT power. The effect of target load (high-low target load) for salient and noisy distractors.The mean is denoted by the horizontal line, the median is denoted by the white dot and interquartile range by the grey vertical bar. C) Target RFT power time courses for high (red) and low (blue) target load relative to cue onset (left) and discrimination target onset (right). Importantly the Target RFT power was elevated for high compared to low target-loads which is consistent with an increase in neuronal excitability in the target with an increased perceptual load. D) Source modelling of the relative baseline target-RFT modulation of high versus low target loads (cue-locked, 500 - 1350ms, shown in B) collapsed over left and right attention, displayed as attention to the left (right hemisphere is contralateral to target). Plot shows top and bottom 1% of the values in the grid-points. E) Correlations between average distractor effect (Target RFT_salient_ - Target RFT_noisy_)/(Target RFT_salient_+Target RFT_noisy_) cue-locked, (500 – 1350 ms after cue onset) and behavioural distractor interference (RT_noisy_ – RT_salient_) of target RFT with respect to low (left) and high (right) target load.

Testing for effects of distractor salience yielded no significant effects, showing that distractor salience does not directly influence input gain of target processing. Instead, it strengthens the existing effect of perceptual load.

##### RFT power associated with the distractor

Analysis of RFT power associated with the *distractor* (Fig. 6A) yielded a significant main effect of ‘distractor salience’ (*F_(1,29)_* = 5.67, *p* = .024, *partial η^2^* = .16), where RFT power overall is stronger for noisy distractors (*M* = 0.67, *SEM* = 0.029) than salient distractors (*M* = 0.64, *SEM* = 0.032). Additionally, there is a significant main effect of ‘target load’ (*F_(1,29)_* = 4.63, *p* = .04, *partial η^2^* = .14), where RFT power is higher when target load is high (*M =* 0.67, *SEM* = 0.027) than when it is low (*M* = 0.64, *SEM =* 0.033), possibly indicating a generic gain increase when target load is high, as this also occurs for the target. Finally, we observed a significant interaction between ‘target load’ and ‘distractor salience’ (*F_(1,29)_* = 15.17, *p* < .001, *partial η^2^* = .35).

**Figure 6.**
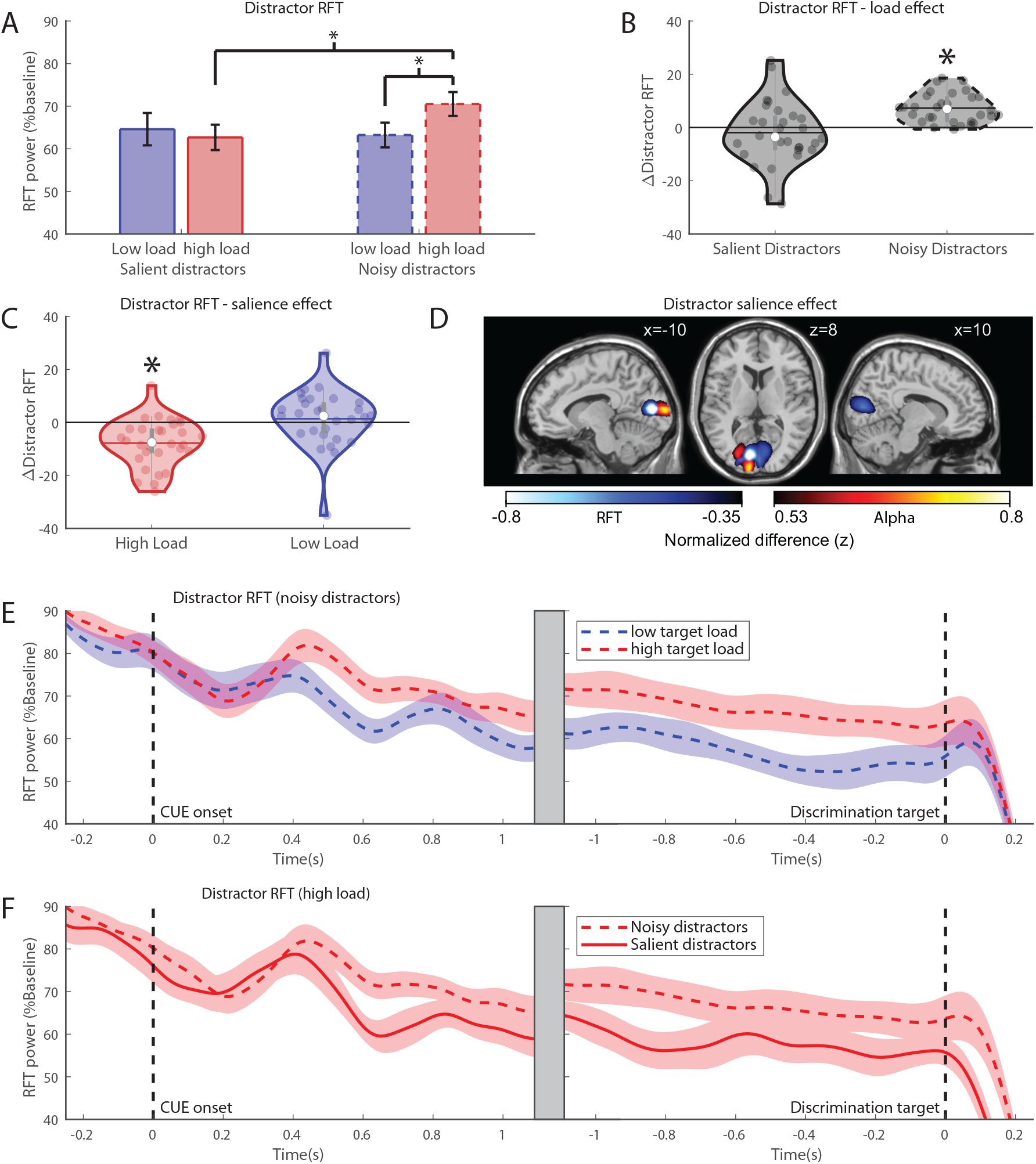
Effects in distractor RFT. A) Average distractor RFT power related to the target for high (red) and low (blue) target load conditions with noisy (dashed) or salient (solid) distractors 500 – 1350ms after cue onset. Error bars denote standard error of the mean (SEM). B) Effect of target load (high-low target load) for salient and noisy distractors.The mean is denoted by the horizontal line, the median is denoted by the white dot and interquartile range by the grey vertical bar. C) The effect of distractor salience (high-low salience) for high and low target load. D) Source modelling of the salient minus noisy distractor difference under high load 0.5-1.1s after cue onset. Cooler colours denote the decrease in RFT power, while warmer colours show the alpha power increase. Contrasts are collapsed over left and right attention and displayed as attention to the left (left hemisphere is contralateral to distractor). Plot shows top and bottom 1% of the values in the grid-points. E) Noisy distractors-RFT power time courses with high and low target load relative to cue onset (left) and discrimination target onset (right) under high target load. F) Distractor RFT power time courses for salient and noisy distractors relative to cue onset (left) and discrimination target onset (right) under high target load.

Testing for effects of target load on distractor RFT revealed a significant effect of target load with noisy distractors (paired samples t-test, *t* = −7.24, *p* < .001, < α = .0125 Bonferroni corrected) but not with salient distractors (paired samples t-test, *t* = .88, *p* = .39 > α = .0125). When distractors are noisy, distractor RFT power increases with higher target load, see figure 6B and 6E for a time-course representation. Here, no significant correlations were found with respect to behavioural distractor interference with both low and high target load (all *p* > 0.5). As can be seen in Fig. 6A, the distractor RFT effects seems to be driven by an increase in RFT power when load is high and distractors noisy. This may be due to a general increase in RFT power with high target load when distractors do not need to be suppressed, as the same effect is present for *target* RFT power with noisy distractors, although this effect does not survive multiple comparisons correction (low load: *M* = 0.82, high load *M* = 0.87, paired samples t-test *t* = −2.37, *p* = .025 uncorrected).

Testing for distractor salience effects revealed significant difference between noisy and salient distractors when the target load was high (paired samples t-test, *t* = 4.80, *p* < .001 < α = .0125), but not when target load was low (paired samples t-test, *t* = 0.71, *p* = .48 > α = .0125), see figure 6C and 6F for a time-course representation. Thus, distractor power is modulated by salience, with reduced RFT power for salient distractors, but only when target load is high. This may indicate that salient distractors are more suppressed than noisy distractors, as also evidenced by increased alpha power for salient distractors in the same condition. Here, no behavioural correlations were observed (*r*_spearman_ = −0.08, *p* = .69). Source modelling indicates that this modulation occurs in central early visual cortex (MNI: x:-10, y:-82, z:10, BA 17), see figure 6D, cooler colors. For comparison, the salience-driven alpha modulation is depicted by the warmer colours and places the strongest source in lateral early visual cortex (BA 17, MNI x:-10, y:-98, z:8). Interestingly, although the modulation of RFT and alpha band power both occur in early visual cortex, the sources do not seem to overlap, where the alpha band modulation is located slightly more lateral and posterior.

## Discussion

Our results show that increasing the perceptual load of a target stimulus increases alpha band power contralateral to distractors, consistent with predictions from perceptual load theory. Importantly, the alpha power increase contralateral to the distractor predicted behavioural measures reflecting the ability to reduce distractor interference, i.e. a failure to increase alpha power contralateral to the distractor resulted in longer reaction times for target discrimination. Rapid frequency tagging (RFT) increased contralateral to targets with high perceptual loads, suggesting increased stimulus gain. Increasing the salience of the distractor stimulus also increased distractor related alpha band power and decreased RFT associated with the distractor, but only when the target load was high. We did not observe an inverse pattern between RFT power and alpha modulations, suggesting that alpha power may not implement gain control. Furthermore, source modelling shows that both RFT and alpha power modulations originate from visual cortex, but sources do not overlap, indicating a distinct cortical origin. Taken together, this indicates that attentional resource allocation driven by the *target* serves to modulate the brain activity associated with both target and distractor processing. This modulation seems to be associated with two complementary mechanisms: a gain increase of the target, as indexed by RFT, and an increase of distractor alpha band power, possibly implementing gating at a later stage.

### Perceptual load results in increased distractor suppression reflected by increased alpha band activity

The results are consistent with the framework of perceptual load theory^2^. In particular, we show that when the perceptual load of the target is increased, alpha band power contralateral to the distractor is increased, as well as RFT related to the target. Thus, the perceptual load of the *target* dictates the resource allocation supporting the task. Behavioural results demonstrate a signficant interference effect on reaction times produced by salient compared to noisy distractors. Importantly, this interference effect interacted with perceptual load: high load eliminated the salient distractor interference effect as predicted by load theory. The electrophysiological findings similarly indicated an increase in alpha band power contralateral to the distractor under high target load which occured when the distractor is salient, but not when it is noisy. Taken together this pattern suggests that the alpha increase under high load results in the suppression of interfering (salient) distractions. The positive correlation, between the strength of modulation of distractor alpha power due to increased target load and the level of behavioural interference caused by the salient (vs. noise-masked) distractors, further supported this conclusion. Given the inhibitory role of alpha band oscillations^8,12,13^, this suggests a reduction or suppression of the resources available for distractor processing under high target load. Thus, resource allocation to the target may occur by reducing resources for distracting stimuli, abolishing interference with high perceptual load. This is in line with a fMRI study by Torralbo *et al.*^38^ measuring BOLD responses under high and low perceptual load. The authors showed increased BOLD responses in early visual cortex to target stimuli with increased load. Importantly, BOLD responses were decreased for irrelevant distractor stimuli. BOLD responses are typically negatively correlated with alpha band power^39–41^. Similarly, using spectroscopy, Bruckmaier *et al*^14^ recently showed increased cellular metabolism for attended target and a reduction for unattended stimuli with increased perceptual load. The current results thus provide evidence that the previously observed resource reduction for the distractor due to a target load increase is implemented by an increase in alpha band power. Similarly, the observed resource increase related to the target may be implemented by an increase in early visual gain, as indexed by the RFT power related to the target in our study.

### The effect of distractor salience

Results show that suppression of distractor stimuli only occurs when these stimuli are actually salient and interfering and task demand (load) is high. This is somewhat expected, given that distractor suppression is thought to be an important part of maintaining task performance^12,42,43^. However, it has recently been questioned to what extent distractor anticipation and processing results in an alpha power increase. Moorselaar & Slagter *et al.*^24^ find no alpha power modulations related to anticipated distractor locations. Likewise, Noonan *et al.*^23^ show that anticipatory suppression of distractors does not occur when the location of an upcoming distractor is given. Although the location of the distractor was not varied in our study beyond the opposite of the target location, we do find evidence for active distractor suppression, through increased alpha band power contralateral to the distractor, dependent on a distractor property (i.e. its salience). This co-occurs with a reduction of distractor-related RFT power. The nature of the distractor stimulus, thus, *does* seem to be important for distractor suppression. Critically, in our study we only find an increase in distractor-alpha power when the perceptual load in target processing is high. This might account for the difference with these^23,24^ studies. In fact, our findings are consistent on this with load theory: the suppression of salient distractors by alpha is only occurring when the target load is high.

### Attentional modulations consist of early gain and downstream gating

Recently, the notion has been put forward that alpha-band oscillations do not impose gain control in early visual regions^32,33^, but rather implement a gating mechanism in downstream visual areas^12,31^. Meanwhile, the RFT response is thought to reflect visual gain^29,30^. These notions would explain why we did not find a consistent inverse pattern of modulations of alpha power and RFT. This inverse pattern was only found when contrasting distractor salience, but not target load. Note however, that an MEG study by Molloy *et al.*^44^ did find reduced alpha power with high perceptual load, both pre- and post- stimulus onset. However, it is unclear whether this was a generic effect or specific to target facilitation. In line with the current findings, Zhigalov & Jensen^31^ recently found no evidence for trial-by-trial correlations between RFT and alpha band power, supporting that gain and alpha band power are independently modulated; as such they are bound to have complementary functions. Also, given the neuronal sources in early visual cortex, the RFT increase likely reflects a visual input gain of the target stimulus, possibly as early as V1. On the other hand, alpha band power contralateral to the distractor increases in visual areas just outside of the calcarine, even extending into fusiform areas. This supports the idea that distractor suppression and target facilitation are driven by different neuronal mechanisms^23^. Taken together, this could indicate a later stage effect preventing further processing of irrelevant items, i.e. a gating mechanism^12^. However, it remains speculative at what stage of visual processing resource allocation critically affects behaviour as effects of perceptual load have been found in a range of visual processing areas^38,45–48^.

### Conclusions

We conclude that the perceptual load of a target stimulus is a key driving factor of attentional modulations. Higher target loads lead to increased suppression of distracting stimuli, reflected by increased distractor-related alpha-band oscillations. Simultaneously, the gain of the target stimulus increases. This results in a reduction of behavioural distractor interference. Attentional modulations are driven by the target load and distractor salience, with target facilitation and distractor suppression working together to achieve a behaviourally favourable outcome and are thus not mutually exclusive. We thus conclude that alpha oscillations reflect distractor suppression with increased perceptual load.

## Materials and Methods

### Participants

N=35 healthy volunteers (25 females, mean age 24.0 years, age range 18-37) participated in the study. Inclusion criteria: English proficient, aged 18-40, right-handed, no history of mental health, normal or corrected to normal vision. Exclusion criteria: history of epilepsy/seizures, medication, MRI contraindications, exceeding scanner weight limits. In line with the criteria set out in our pre-registration (https://osf.io/ha4vw/, 1/10/2018), five subjects were not entered into the main analysis due to failure of one or more of the criteria: Poor data quality due to MEG artifacts and/or subject (eye) movement, resulting in a rejection of more than 1/3 of the trials (4), marginal frequency tagging response (1). The study was approved by the science, technology, engineering and mathematics ethical review committee of the University of Birmingham. All subjects signed an informed consent before participation and were paid 15GBP per hour.

### Stimuli and Procedure

During MEG acquisition, subjects performed a cued change detection task (Fig. 1A). To allow more space in the bottom half of the screen, the fixation was shifted upwards by 20% of the screen. While fixating, subjects were presented with two face stimuli (circular, 8° visual angle diameter, 7° eccentricity from fixation to stimulus centre) in the left and right lower quadrant of the screen. After 1000ms, a directional cue (presented for 350ms) appeared at the location of the fixation spot (0.7° height and width) to instruct which hemifield is to be attended. After a variable 1000-2000ms interval, a change occurred in the eyes of both face stimuli (the pupil shifted 25% of the eye width horizontally such that the gaze of the presented face stimuli shifted left or right). The direction of gaze change was varied randomly for either of the two presented faces. The faces with the new gaze were shown for 150ms and subsequently disappeared. The subject’s task was to identify the change in the cued face and respond as quickly as possible by button press using either the left (leftward gaze) or right (rightward gaze) index finger on a set of MEG compatible button boxes (NAtA technologies, Coquitlam, BC, Canada).

Increasing the perceptual target load was done by masking the target stimulus with noise^46^, while reducing the saliency of the distractor stimulus was done by masking it with noise (noise mask as for target), see figure 1C. To maintain the same distribution of pixel intensities, and thus frequency tagging power, the noise mask was created by shuffling 50% of the face pixels to a random location. These manipulations allowed for modulating the target load as well as distractor saliency. Eight different face identities were used. The same faces were used in the left and right hemifield for every trial, although the left and right face were mirror symmetric from the fixation point, and the 8 identities were randomised across trials. Stimuli were presented using a Windows 10 computer running Matlab R2017a and Psychtoolbox 3^49^. Stimuli were projected using a VPixx PROPixx projector (VPixx technologies, Saint-Bruno, Canada) in Quad RGB mode (1440Hz) with an effective resolution of 960×540 pixels. A projection screen was placed 148cm from the participant, spanning 71cm, resulting in a visual angle of 25.6°. The left and right face stimuli were ‘frequency tagged’, i.e., the luminance of the white parts of the face stimuli oscillated sinusoidally at a fixed frequency, see figure 1B. The tagging frequencies used were 63Hz and 70Hz, which are above the flicker fusion threshold, and the ‘flicker’ was thus effectively invisible to the subject and, importantly, did not create an imbalance in stimulus salience.

### Data acquisition

The ongoing MEG data were recorded using the TRIUX™ system from MEGIN (MEGIN, Stockholm, Sweden) with subjects in upright position. This system has 102 magnetometers and 204 planer gradiometers. These are placed at 102 locations, each having one magnetometer and a set of two orthogonal gradiometers. The vertical EOG, eye tracker data (EyeLink 1000, SR research Ltd., Ottawa, Canada) and polhemus (Polhemus, Colchester, USA) scalp surface data were acquired together with the MEG data. For accurate source localisation, an anatomical T1 MRI scan was acquired for each participant (Philips Achieva 3T, 23 subjects, Siemens Magnetom Prisma 3T, 10 subjects) if not available from a previous source. The MEG data was lowpass filtered at 330 Hz, sampled at 1000Hz and stored for offline analysis. Subjects completed 2 blocks of 256 trials, 512 trials total, lasting approximately 45 minutes excluding short breaks or 60 minutes including breaks.

### Data analysis

*Reaction times* (RTs) were obtained from the subjects’ button box responses. RTs shorter than 100ms or longer than 1000ms were excluded, as well as trials with eye movements (inspected using EyeLink eye tracker data) larger than 3°. Behavioural data from rejected MEG data trials was rejected to keep consistency between the behavioural and electrophysiological data. All data were sorted by condition (target load condition x distractor salience) and averaged per participant. To assess the effect of distractor interference per target load condition, the distractor interference index (DI) was calculated as:

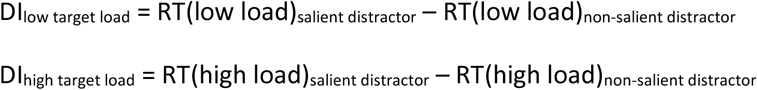

*MEG data* were analysed using the Fieldtrip software package^50^ and custom scripts in the Matlab environment. The analysis pipeline scripts are available at https://github.com/tgutteling/Perceptual_Load_Experiment. In short, Raw MEG sensor data were demeaned and high-pass filtered at 1Hz. For the cue-locked analysis, data were segmented in 3.3s epochs, with 2.3s prior and 1s after cue offset. For discrimination onset-locked analysis, data were segmented 4s epochs, with 3s prior and 1s after onset of the discrimination target (eye movement of the face stimuli). Independent component analysis (ICA), using the ‘runica’ algorithm was used to ’project out’ artefacts after removal of trials with gross incidental artifacts (e.g. jumps) and noisy sensors (sustained gross artifacts) manually. Component removal was restricted to ocular, cardiac and muscle artefacts. After ICA clean up, residual artifacts were identified by visual inspection. Trials containing clear residual artifacts and deviations from fixation larger than 3° (inspected using EyeLink eye tracker data) and eye blinks (inspected using vertical EOG) in the critical −1s – 1s interval relative to the cue offset, or the −1.5s – 0s interval relative to the discrimination target onset were rejected. On average 13.7% (*SD* 8.0) of the trials were rejected. Removed sensors were interpolated using a weighted neighbour estimate.

For *frequency domain analysis*, data were analysed separately for the high frequency range (50-100Hz) and the low frequency range. For the high frequencies, power modulations were estimated using a fixed 500ms sliding time-window, with steps of 1ms and a 4s zero padding to obtain integer frequency bins, resulting in a frequency smoothing of ~3Hz. A single taper approach was taken to avoid spectral smearing of the narrow frequency of interest (the tagging frequencies). Power was calculated using FFT after the application for a 500 ms Hanning taper. For the lower frequencies, a 3 cycle time-window (e.g. 300 ms for 10 Hz) was used with 1ms steps and next power of 2 zero padding for efficiency, resulting in a frequency smoothing of ~4-6.5Hz in the alpha range (8-13Hz). After frequency domain analysis, the planar bidirectional gradiometers were summed. From the resultant spectral estimates, *power time courses* were extracted for every subject and condition from a sensor cluster. For both alpha range and RFT time courses, the four sensors that showed the strongest attentional modulation at group level (i.e. attention left minus right after cue offset) were chosen. Time courses were normalised using a relative baseline based on the pre-cue baseline 350ms after stimulus onset until 400ms before cue offset (600ms total) per hemifield and frequency.

To test for the behavioural effect of perceptual load, *behavioural data* (reaction times) were entered into an ANOVA with factors ‘target load’ and ‘distractor salience’. The same analyses were conducted for the mean alpha band power extracted from the sensors contralateral to targets and distractors, 500ms after cue onset until the earliest time at which the discrimination could appear (1350ms after cue onset). This time was chosen to avoid contamination from evoked responses from the cue.

To establish a link between behaviour and electrophysiology, *correlations* between the behavioural distractor interference (DI) and RFT/alpha power were calculated using the same time window as the main ANOVA analysis (500-1350ms after cue onset), across subjects, using Spearman rank correlation.

*Source localisation* was performed to identify the regions producing the power modulations. In each subject, the scalp surface obtained from the Polhemus digitisation was combined with the MRI T1 scan to obtain an individual realistic single-shell head model. These were subsequently used for source estimation using the dynamical imaging of coherent sources (DICS) beamformer approach^51^. The brain volume was divided into a 5mm grid. Oscillatory activity of interest was estimated using DPSS tapers with a frequency smoothing of 3Hz and a centre frequency of 10.5Hz for the alpha band and 63/70Hz with 2Hz smoothing for RFT. All source analyses focused on the same time window as the main ANOVA: 500-1350ms after cue offset. A common filter was estimated by pooling across all conditions that were contrasted. For alpha band sources, contrasts were created by subtracting the power in the time window of interest for both conditions and normalising by the sum of power for these conditions. For RFT sources, a contrast was first made with the same pre-cue baseline period as in the time-course analysis, 650ms after stimulus onset until 400ms before cue offset. Contrasts of interest were subsequently created form the baselined source estimates. For visualisation, contrasts for attention right were mirrored along the y-axis, normalised to non-flipped MNI space and averaged with the attention left condition.

